# Deep Directed Evolution of Solid Binding Peptides for Quantitative Big-data Generation

**DOI:** 10.1101/2021.01.26.428348

**Authors:** Deniz T. Yucesoy, Siddharth S. Rath, Jacob L. Rodriguez, Jonathan Francis-Landau, Oliver Nakano-Baker, Mehmet Sarikaya

## Abstract

Proteins have evolved over millions of years to mediate and carry-out biological processes efficiently. Directed evolution approaches have been used to genetically engineer proteins with desirable functions such as catalysis, mineralization, and target-specific binding. Next-generation sequencing technology offers the capability to discover a massive combinatorial sequence space that is costly to sample experimentally through traditional approaches. Since the permutation space of protein sequence is virtually infinite, and evolution dynamics are poorly understood, experimental verifications have been limited. Recently, machine-learning approaches have been introduced to guide the evolution process that facilitates a deeper and denser search of the sequence-space. Despite these developments, however, frequently used high-fidelity models depend on massive amounts of properly labeled quality data, which so far has been largely lacking in the literature. Here, we provide a preliminary high-throughput peptide-selection protocol with functional scoring to enhance the quality of the data. Solid binding dodecapeptides have been selected against molybdenum disulfide substrate, a two-dimensional atomically thick semiconductor solid. The survival rate of the phage-clones, upon successively stringent washes, quantifies the binding affinity of the peptides onto the solid material. The method suggested here provides a fast generation of preliminary data-pool with ∼2 million unique peptides with 12 amino-acids per sequence by avoiding amplification. Our results demonstrate the importance of data-cleaning and proper conditioning of massive datasets in guiding experiments iteratively. The established extensive groundwork here provides unique opportunities to further iterate and modify the technique to suit a wide variety of needs and generate various peptide and protein datasets. Prospective statistical models developed on the datasets to efficiently explore the sequence-function space will guide towards the intelligent design of proteins and peptides through deep directed evolution. Technological applications of the future based on the peptide-single layer solid based bio/nano soft interfaces, such as biosensors, bioelectronics, and logic devices, is expected to benefit from the solid binding peptide dataset alone. Furthermore, protocols described herein will also benefit efforts in medical applications, such as vaccine development, that could significantly accelerate a global response to future pandemics.

## Introduction

Directed Evolution libraries such as phage display, cell-surface display, mRNA display, and yeast display, are powerful tools for the identification of peptides and proteins, including enzymes and antibodies, with an affinity for a specific target such as antigens, drugs, organic molecules, and inorganic materials.^1-6^ Over the years, the authors and others have successfully applied the tools, particularly M13 phage display and bacterial cell surface display (FLITRX), to study peptide-solid interactions for a myriad of bio-nanotechnological and biomedical applications.^2^ The link between phenotype and genotype of organisms is the common feature in all combinatorial display techniques where the randomized (variant) peptide sequences are displayed as fusion partners with different surface proteins.^2,6-8^ The authors are one of the pioneers of adapting DE techniques to select peptides with affinities to metals, ceramics, semiconductors, and minerals with about 5000+ Solid Binding Peptides (SBP) specific to 25+ different materials.^2,9-11^ SBPs that are successful as fundamental building blocks as molecular linkers, erectors, and assemblers in bio-nanotechnology implementations, are termed Genetically Engineered Peptides for Inorganics (GEPI). Using a unique representation of data and conventional bioinformatics tools, the authors also discovered tiny synthesizers from selected peptides that catalyze solid materials synthesis, e.g., gold, silica, and hydroxyapatite, from ionic precursors.^11,12^ The phage display selection procedure for biomacromolecules is well-established. But the technique can achieve even further improvement in selectivity by integrating simple modifications into the biopanning protocol (e.g., counter-selection step, material specificity testing, etc.) to isolate SBPs with high affinity and specificity to the desired material.^2,7,13,14^ Detailed procedures, therefore, are developed for a particular inorganic material in the powder, thin-film, or in single crystal forms and are demonstrated in numerous publications.^9,12,15-17^

The selected number of peptides has conventionally been small in the literature, up to a few tens of SBPs.^9,15^ The authors ensure that in any SBP-selection, the number of GEPI identified for a given material (either used as a single crystal or a powdered form) are at least 35 (with an average of 50, max 96 peptides). Despite well-established phage display selection procedures and subsequent improvements towards increasing the winner peptides’ potential, the methods are still far from leveraging the huge combinatorial potential since the diversity of the libraries is in the order of 10^9^ variants.^2,7,15,18,19^ The drawback is primarily due to the limited scalability of clone selection techniques and characterization. Moreover, in low-throughput Sanger sequencing workflows traditionally used, the number of peptides isolated in a reasonable time frame after several rounds of the biopanning process is limited to 10-100 clones. Compared to the vast diversity of the naive library (∼10^9^ unique sequences), the conventional peptide-selection approaches are disappointingly low-throughput (10-100 peptides) and provide a minimal perspective (<0.00001%) of the complete variant (sequence) space.

Such a small sample size is, therefore, not only prone to bias from nonspecific, preferentially amplifying false-positive hits but also leads to omitting a large number of promising candidates.^2,7,15,18^ Traditionally, peptides enriched after several rounds of selection are identified by DNA sequencing of the inserts of a limited number (tens to hundreds) of clones. Depending on the sequence diversity remaining in the library after selection, the analysis of a manually chosen limited number of clones does not necessarily result in discovering the most promising candidates. Moreover, phage display screenings are notorious for identifying false-positive hits’, e.g., parasitic sequences”.^14,20,21^ Such sequences emerge for two crucial reasons: (a) Binding to materials used during the selection other than the desired substrate (such as plastics or albumin), and (b) Propagation advantages.^4^ A well-known example in the latter category is the greatly accelerated propagation of phages displaying the HAIYPRH peptide in the Ph.D.-7TM library due to a mutation in the Shine-Dalgarno box of the phage protein gIIp in this clone.^5^ This peptide has been identified in at least 13 independent biopanning experiments.^4^ Several web-based tools aid in identifying potential false positives, viz., PepBank can be used to search for peptides already published in other experiments,^22^ while SAROTUP searches for peptides binding to unintended materials.^23^

For the accelerated design of soft interfaces, device development, and deployment, leveraging statistical inference and Machine Learning (ML) tools is necessary. Developing ML models is critical not only for a practical exploration of the sequence space but also to make the experimental procedure more efficient and targeted for a variety of bio-nanotechnological applications. To generate large SBP/GEPI data sets to develop an ML algorithm to predictively design peptides, the authors incorporated Next-Generation Sequencing (NGS) tools and screened millions of peptide sequences in one shot. Such a method allows a more comprehensive look at the phage display library’s sequence-space, enabling a more practical and higher resolution characterization of the library.^24-26^ However, the lack of a high-throughput fluorescent-microscopy (FM) or spectroscopy-based^27^ end-point binding characterization method prevents the characterization of binding affinities of SBPs, which is necessary to identify GEPI and for labeling of the massive training data in ML algorithms. In the absence of conventional binding-affinity information, here we report a high-throughput peptide-selection protocol with functional scoring to characterize the peptide affinity to the surface as a function of its count number (survival probability) and ML-based data analysis models to validate the scoring.^28,29^

Machine learning aided directed evolution is more effective at designing SBPs than wet-lab guided DE alone.^30-35^ ML models are paramount in generating and analyzing sequence libraries more targeted towards a specific function, such as the device behavior in bio/nano interfaces. An ML integrated DE, and NGS platform necessitates generating a custom phage library from the predicted sequences that can be screened in a high-throughput manner to validate and benchmark the predictions. The implications of such an integrated experimental and mathematical platform are varied but suffice to mention that it is imperative to meet the demands of modern scientific and application-focused research at the bio/nano interfaces. Identifying and analyzing millions of SBPs in a single shot will decipher underlying mechanisms of evolution towards specific bio/nano interaction. It will also develop superior peptide prediction platforms to design novel peptides with known functions towards various technological and medical implementations (Fig 1).

**Fig 1.**
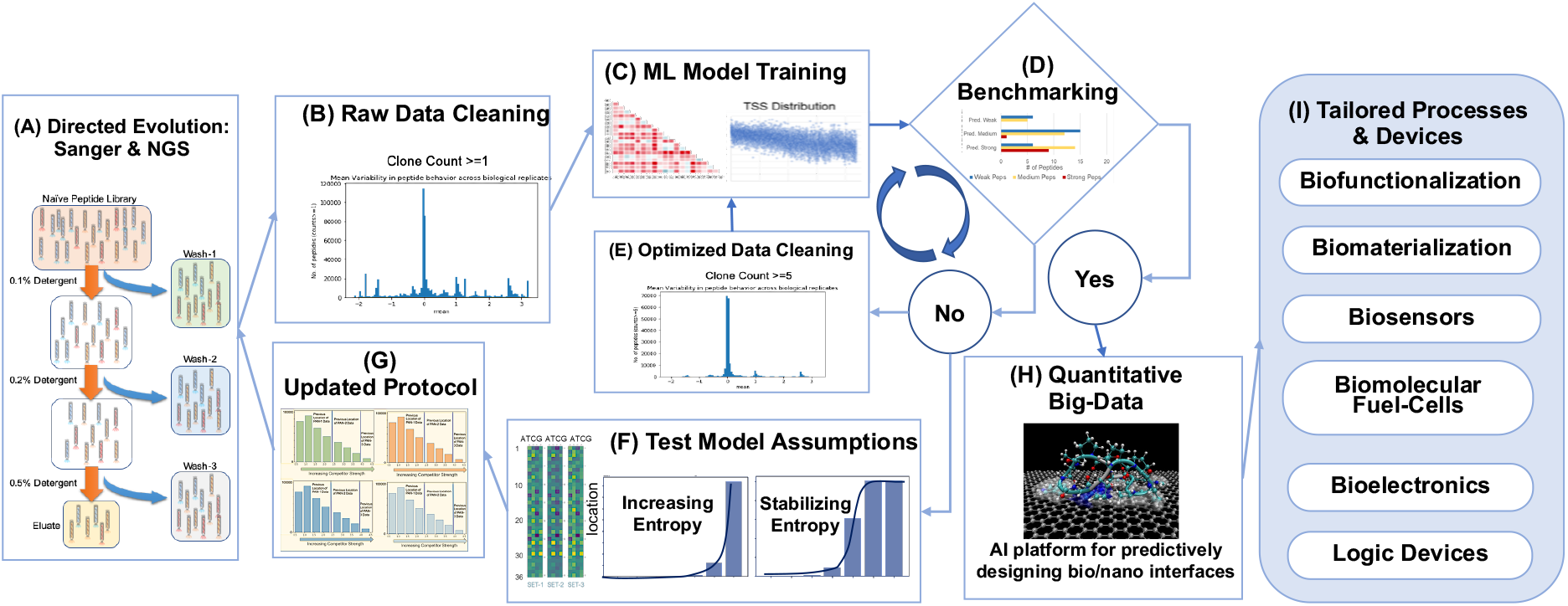
Overall schematic of the iterative deep directed evolution process for smart peptide selection. (A) through (H) enumerate the steps in the procedure. Conventional DE is used, with amplification for creating benchmark datasets while NGS is used for big-data generation with later iterations guided by ML/AI models. (I) Deep directed evolution enables faster design of functional bio/nano materials.

## Materials & Methods

### Deep Directed Evolution for Big-Data Generation

#### Combinatorial Mutagenesis and Biopanning

The 12-mer Phage display (PhD) library with an estimated diversity of 1.7×109 different clones of *M13* bacteriophage is used to select peptide sequences fused to the minor coat protein (pIII) against MoS_2_ flakes (∼300 mesh, 99%, 10129, Alfa Aesar). Before the screening process, MoS_2_ flakes are cleaned by sonication sequentially in isopropyl alcohol and de-ionized water, then dried under vacuum. For high-throughput isolation and screening of MoS_2_ binding clones, 5 mg of MoS_2_flakes are dispersed in 1 mL of potassium phosphate/sodium carbonate buffer (PC, 55 mM KH2PO4, 45 mM Na2CO3, and 200 mM NaCl, pH 7.4), containing 0.02% Tween 20 detergent (Merck, Whitehouse Station, NJ, USA). 10 µL of the PhD library (1011 pfu, New England Biolabs) is added onto dispersed flakes and incubated for 3 hours on a rotator at room temperature and then washed twice before overnight incubation. After incubation, to remove the nonspecifically- or weakly-bound phages, the MoS_2_ flakes are washed with PC buffer with increasing detergent concentrations as follows; 0.1% (v/v; Wash Round 1), 0.2% (Wash Round 2), 0.5% (Wash Round 3), at pH: 7.4. The remaining ‘strong’ bound phages are then eluted from the surface in a stepwise manner by applying an elution buffer consisting of 0.2 M Glycine-HCl pH 2.2 (Sigma Aldrich, St. Louis, MO) for 15 min. The washed off, and eluted phages are then transferred to a fresh tube and neutralized. They are labeled as Wash-1, Wash-2, Wash-3, and Eluate. Each phage pool is then purified via PEG/NaCl precipitation and resuspended in de-ionized water. The overall selection procedure is performed in three biological replicates and labeled as Set-1, Set-2, and Set-3 and two technical replicates that are sequenced separately but subsequently combined during data analysis.

#### DNA Isolation and Next-Generation Sequencing

DNA amplicons are prepared as previously described with slight modifications.^36^ Single-stranded DNA (ssDNA) is isolated from wash and eluate phage pools using QIAprep Spin M13 Kit (QIAGEN). The sequencing library is prepared by amplifying the 36 bp peptide-coding variable region. Q5 polymerase (New England Biolabs) is used to amplify the target region with forward and reverse primer sequences given below:

Forward: CCGCGTGATTACGAGTCGCAAGCTGATAAACCGATACAATTAAAG Reverse: GGGTTAGCAAGTGGCAGCCTACGTTAGTAAATGAATTTTCTGTATGGG. Illumina sequencing adapters containing the p5 and p7 index sequences are attached using a second PCR. The purified PCR products from each phage pool are loaded in duplicates (technical replicates) on the sequencer plate and sequenced on the Illumina NextSeq platform. The Next-Generation Sequencing (NGS) platform yielded 288M DNA sequences in total. About 1.3M and 1.1M individual peptides were obtained (replicate-1) in wash-1 and eluate pools, respectively, where ∼433124 of the sequences existed in both pools. After combining with the other replicates, upon subsequent translation into amino acid sequences, more than 2 million unique peptides were identified with varying copy numbers that survived on MoS_2_ with successively stringent washes. The survival probabilities are then computed as described in later sections to provide a relative label for each unique sequence.

A conventional Directed Evolution procedure, outlined in the supplementary document (S1), is used for benchmark data generation. The overall process is visualized in Fig 2. We choose survival probability as a label for peptides obtained from the high throughput deep directed evolution experiment but use spectrophotometry or Fluorescence Microscopy assays to label benchmark peptides. Readers can also find the characterization procedure for benchmark peptides in the supplementary documents (S1 and S1 Fig). The discrepancy opens an avenue for transfer learning and metric learning for prospective statistical models. Survival probability with subsequent washes is not yet an established industry standard for binding affinity measurement, and as such, researchers should not label benchmark data with such a metric. However, future ML or AI models could establish survival probability as a reasonable measure for affinity prediction, thereby removing low-throughput characterization bottlenecks and opening up avenues for even more high throughput data collection for a myriad of applications. However, data-scientists must clean massive data appropriately before using it in ML/AI models, so we describe our current data cleaning process in the next section.

**Fig 2.**
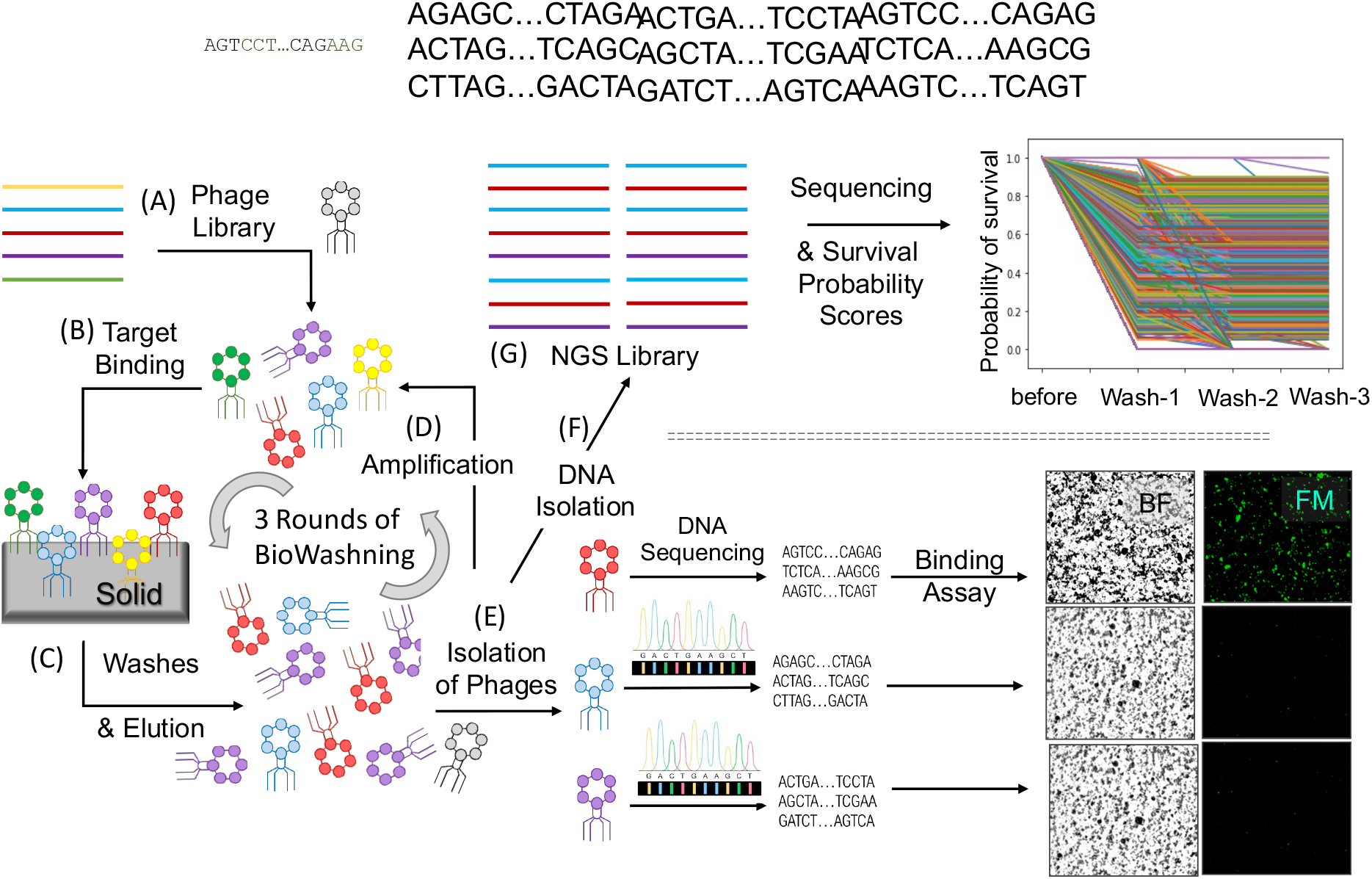
Detail Schematic of the Deep Directed Evolution process. (A) Variant library is created by cloning random DNA fragments into M13 phage and (B) Exposed onto target solid material which is followed by (C) Series of washing and (D) Elution/amplification steps. For benchmark set, this is followed by (E), (F) isolation and sequencing of the phages (before binding affinity characterization) from the colonies while for deep selection, it is followed by (G) next generation sequencing.

### Preliminary Data Cleaning

Traditional peptide selection and characterization provide a quantitative analysis of the peptide binding as relative surface coverage or phage-retention, obtained from fluorescence microscopy or spectrophotometric analysis. The NGS data, however, does not provide such information explicitly.^37,38^ It generates a massive amount of sequence data and their copy number in the collected samples, prior and posterior to each round of wash. To quickly characterize the sequences selected through multiple washing steps, one needs to process the raw sequences and their abundances to establish a correlation between the survival probability of each peptide through the successive washing steps and their binding affinity to the surface. Cleaning of the dataset obtained would result in more robustness in the calculation of survival rates for each sequence and impart more clarity as to which mathematical models should be applied to the data. In the following paragraph, we describe the cleaning steps in detail, including consolidation of DNA datasets, translation into peptides, before obtaining and applying thresholds for filtering the dataset from a raw to a cleaned set, ready for ML model application.

We convert the raw data obtained as FASTQ files from the NGS experiment to TSV format before the DNA sequences’ computational translation into peptide sequences. We then get the raw, but assembled DNA reads where the sequence is labeled with its abundance (termed variously throughout the document as population, count, or copy number) in the sample. We label each sample by its biological replicate (Set-1, 2, or 3), the washing step (wash-1, wash-2, wash-3, or eluate), and its technical replicate (a or b). Overall there are about 288M non-unique sequences with significant overlaps between the replicates.

In the next step, we combine the technical replicates and sum the populations of overlapping sequences, followed by concatenating the different datasets for each wash. As a result, we obtain consolidated DNA datasets for each biological replicate. The first column lists all the sequences, and the following four columns list their populations (combined technical replicate populations) in the various washes. We end up with ∼55M unique sequences, with individual counts in washes and replicates transformed by a logarithm of base 2. The logarithm transform is crucial because every DNA sequence with n copies ends up with at least 2n copies in a usual PCR amplification process. After that, we convert the DNA sequences into Amino Acid sequences. If any two unique DNA sequences encode the same peptide, we sum their populations by wash label (wash-1, 2, 3, or Eluate). We impose restrictions on the populations, where, if a DNA sequence occurs three times or less in total, we do not include it in the translated peptide dataset. Therefore, we end up with about 2 million unique peptide sequences after the preliminary cleaning process discussed above.

## Results & Discussions

### Data exploration and conditioning

After the preliminary cleaning and analysis, with >97% overlaps among the biological replicates, we calculate the survival probabilities for each peptide sequence per biological replicate. Per replicate, we sum the copy numbers of each unique peptide over all the washes and eluate to obtain a total ‘starting’ population. Based on the total population, we calculate the probability of survival as shown in the equations in Fig 3. Fig 3 also shows the survival probability trends of a sample of sequences over the washing steps and eluate, for a biological replicate. Finally, we multiply the survival rates together, and stagger them as shown in Fig 3, to obtain a single functional score: ‘survival affinity’, as a measure of overall phage retention on the surface. We then categorize the peptides as strong, medium, or weak binders based on the distribution of survival affinities. We assume that survival affinity should correlate with the actual phage retention or surface coverage values obtained for the Sanger-based benchmark set (see supplementary S1).

**Fig 3.**
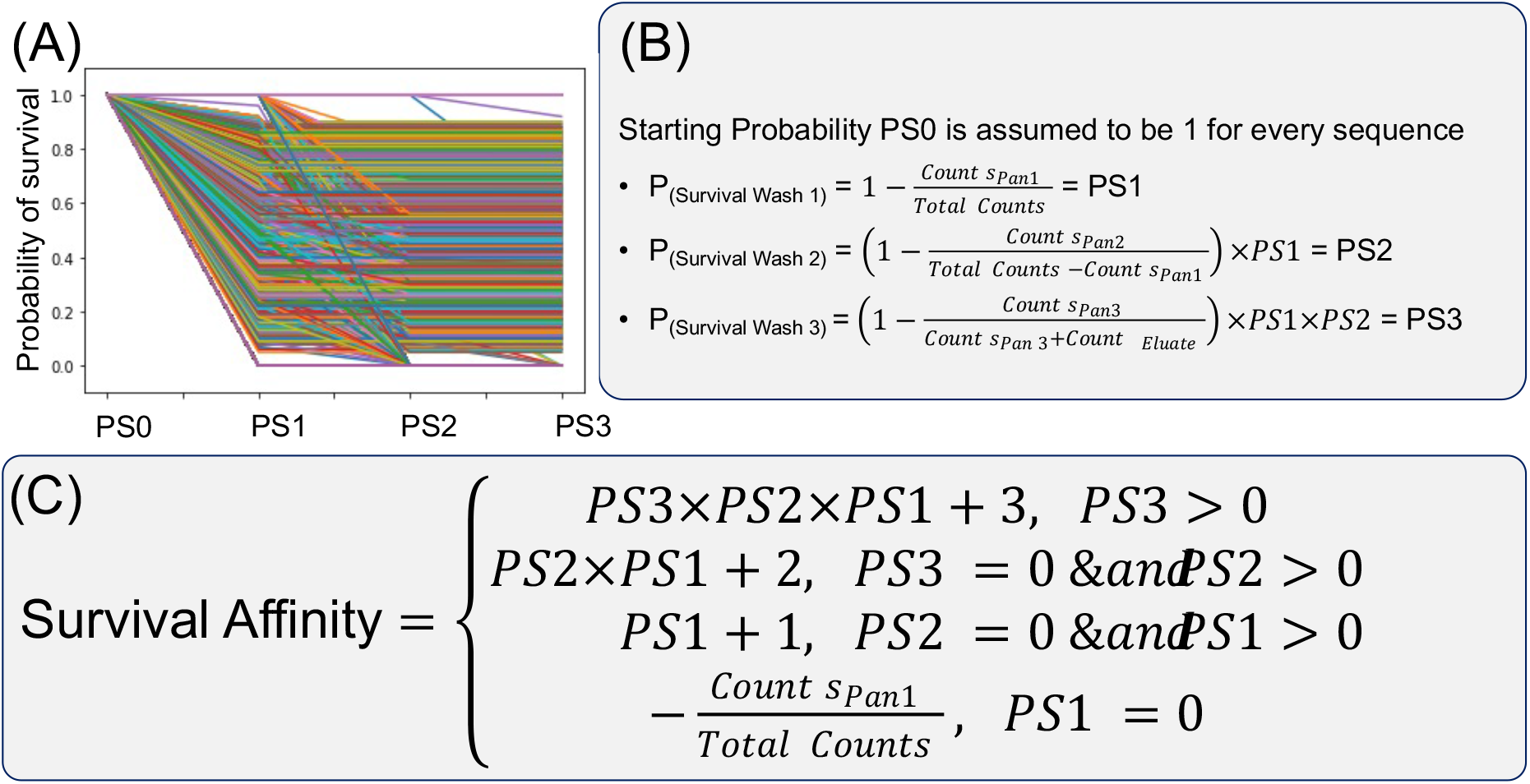
Functional Scoring for labeling massive sequence data. (A) Survival probability trends for a random sample of sequences. PS0, PS1, PS2, and PS3 are the probability of survival (probability of finding a sequence) before washing, after wash-1, after wash-2, and after wash-3 respectively, obtained from equations in (B). (C) Obtaining a single ‘survival affinity’ metric, the functional score, by multiplying and staggering the survival probabilities as shown.

After the cleaning process, we condition the data to make it ready for subsequent data analysis, such as custom statistical algorithms (discussed in later sections) and similarity analysis. We also obtain the ‘center of abundance-mass’ for each sequence, i.e., the sum of the product of population in the wash and wash index (1 through 4; 4 being the eluate) divided by total population as shown. The new metric is important because many sequences are represented in all the four washes including the eluate. While the survival affinity measure follows a Poisson-like distribution with multiple outliers, the center-of-abundance-mass is Gaussian distributed, also with numerous outliers. Due to the high number of outliers, i.e., significant outlier modes, we impose two other cleaning criteria to condition the dataset properly. The first criterion is that the same peptide’s survival trends in three different biological replicates must be about the same. Therefore, the mean difference in survival affinity between biological replicates for the same peptide must be zero. However, the mean difference among sets is nonzero with multiple modes and noisy distribution (Fig 4). We, therefore, impose a minimum copy number threshold (Fig 4). After the count restriction, we keep all sequences that behave similarly in at least two out of three biological replicates (Fig 4) and exclude the rest. As a result, the final well-cleaned and conditioned dataset contains upwards of a hundred thousand unique sequences in total, with their count numbers in all the washes and eluate, their survival affinities, and center-of-abundance-mass values.

**Fig 4.**
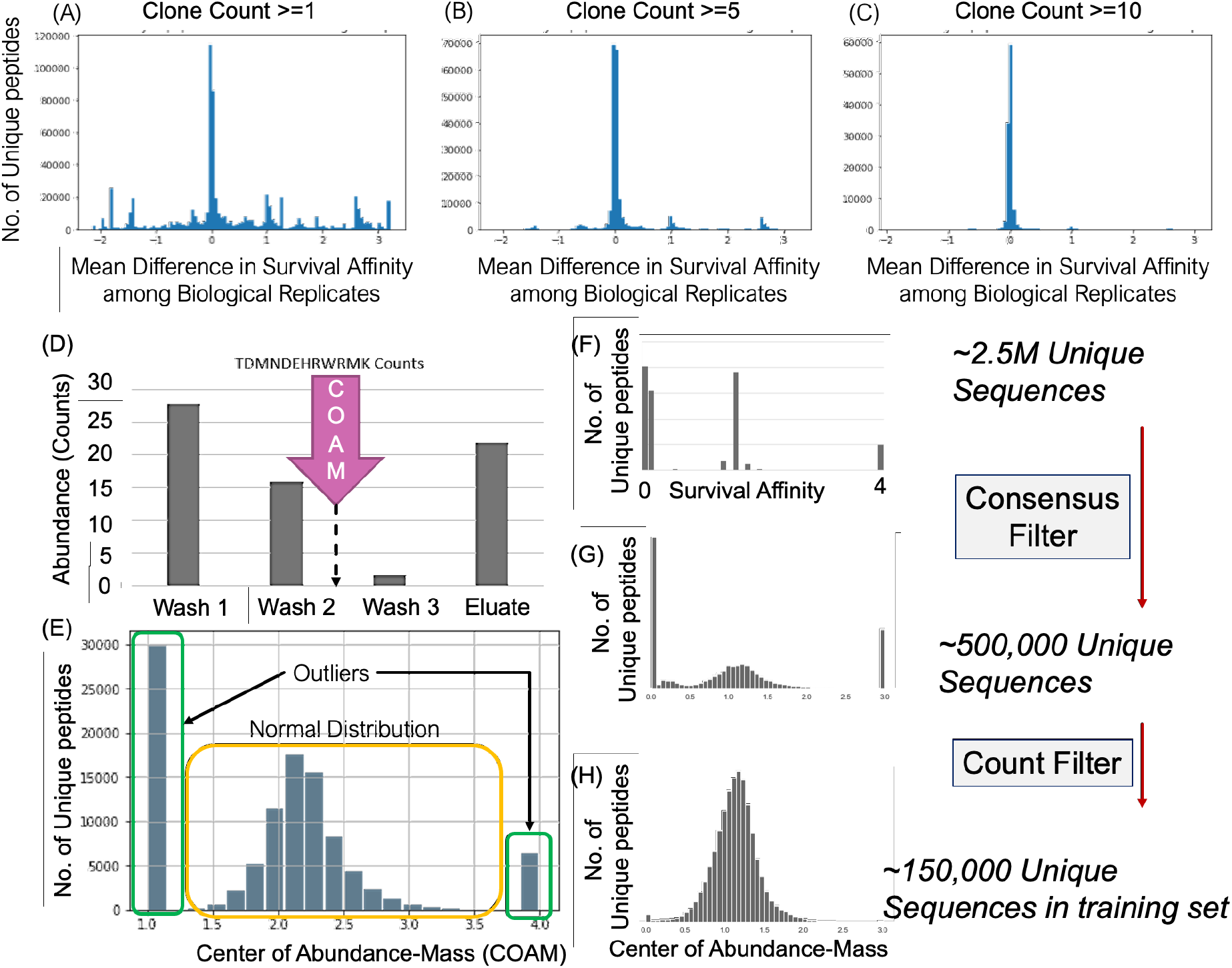
Data Cleaning, Conditioning and the ‘Center of Abundance Mass’ metric. (A)-(C) Upon imposing a minimum count threshold over the sequences, we see a marked decrease in noise and outliers in the mean difference in survival behavior for the same sequence in the three biological replicates. A mean difference of zero implies that the sequences behave in the same manner in all three replicates (Set1, Set2 and Set3). (D),(E) Most sequences appear in all four pools (wash1-3, Eluate). A Center of Abundance Mass (COAM) value for such a sequence is shown to be continuous variable falling anywhere between 1 and 4, with massive outlier modes at the extreme values. (F) We only consider peptides that display similar survival affinities in 2 out of 3 biological replicates. (G) We replace survival-affinity with the COAM value. (H) We impose the clone count threshold, and finally end up with about 150,000 unique sequences in the final dataset.

We split the now cleaned and conditioned dataset into two parts. We set aside 90% of the sequences to be the training set for predictive modeling, while the remaining 10% constitute the test-set for the same. The benchmark dataset is the 96 Sanger-based sequences. In earlier work, authors show that the binding affinity of a peptide of interest is proportional to the sum of the Global Alignment scores to a group of highly functional peptides.^39,40^ We expand upon the concept and apply several custom preliminary bioinformatics and machine learning approaches to the data described in the supplementary information (S2 Fig and S2). Surprisingly, even increasingly complex models (failed to reconcile predicted survival-affinity trends satisfactorily with observed trends in the test set and phage-retention trends in the benchmark-set.

The various models described in the supplementary (S2) that we apply to the data had an underlying assumption that (a) the eluates are the strong binders and that (b) strong binders have the highest self-similarity in their sequences. We analyze the combinatorial diversity encapsulated in the washes through the means of information entropy.^41^ We find that the eluate diversity is higher than all the other washes before; therefore, either (i) the eluate does not contain just the strongest binders, or (ii) strong binders do not have higher self-similarity than weak binders. However, previous studies can rule out the second option as they have conclusively proven that strong binders do indeed have high self-similarity^42,43^. Therefore, the eluted peptides are not just the strongest binding set. Indeed, peptides from previous washes (wash-1 through 3) also exist in the eluate, although in far fewer numbers. Therefore, the eluate contains the spectrum of strong MoS_2_ binding peptides with a mechanism that was attacked by the tween detergent and includes the entire range of binding strengths through various other mechanisms that tween detergent did not attack at all.

### Analysis of sequence diversity to design future experiments

Next, we investigate the combinatorial coverage via information entropy analysis, thereby exploring the relative diversity of the DNA sequence space captured for each wash and the eluate. We weight the sequences by their copy numbers to keep the analysis unbiased (i.e., use conditional probabilities instead of regular ones). Shannon’s entropy, S_info_ is a concept taken from statistical physics but repurposed to describe information content by Claude Shannon^41^ and is mathematically expressed in equation (1):

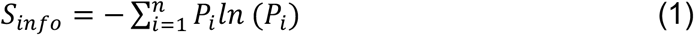

Where P_i_ is the probability of the i^th^ state of a random variable that can take multiple states from 1 to n with varying probabilities. Applied to the case of DNA sequences, the finite number of states, i, that each location on the sequence can take, are either A, C, T or G. In the current dataset, there exist 36-length DNA sequences that code for 12-amino acid long peptides. Therefore, for any given sample of sequences, a matrix with 36 rows (locations on the sequence) and 4 columns (nucleotides) can be created where the sum is 1 for each row. Using equation (1) on samples in the washes, and conditioning on the same probabilities for samples in the input library (the input library is estimated by pooling the sequences obtained from all the Washes, including eluates together).

The total sample information entropy measures the combinatorial coverage of sequences in the sample with n sequences. We estimate maximum entropy for the DNA sequences to be 49.90 (arbitrary units of ‘nats’). Applying the above formalism to the washes, the authors observed that the combinatorial coverage of wash-1 is greater than wash-2. Similarly, the combinatorial coverage of wash-2 is greater than wash-3 (Fig 5).

**Fig 5.**
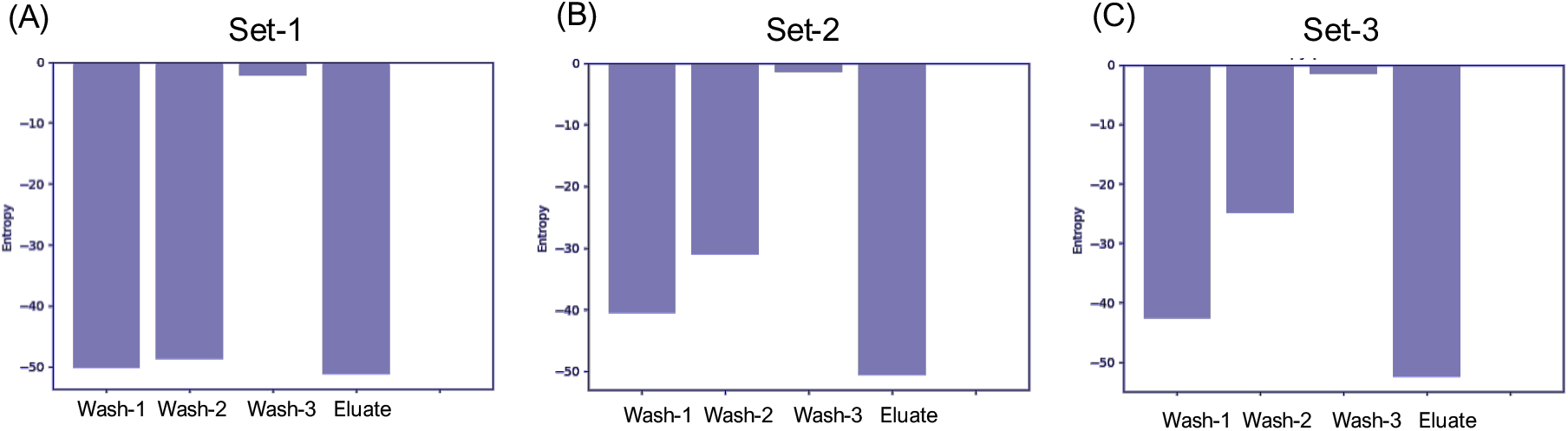
Information Entropy measures combinatorial diversity in the washes and Eluate. (A)-(C) A greater negative value indicates more Information entropy (arbitrary units), more diversity in the pool (wash1-3, or eluate) across all biological replicates. As can be seen, the eluate has similar or higher diversity than all the washes, implying that the eluted sequences are more dissimilar to each other than those in the washes. We therefore cannot assume that the eluted phages are the strongest binders. Simply that the tween detergent used in the washes did not disrupt the mechanism of interaction between the eluted phages and the substrate. We thus propose a future experiment with washing performed with solutions that target specific mechanisms of substrate binding. A simple denaturation may not be enough to disrupt the interactions at the bio/nano interfaces.

Results in Fig 5 tell us that, indeed, sequences considered weak binders, i.e., sequences washed away in the first wash, are more diverse than the successive washes and have little correlation among themselves, as expected. It comes as no surprise that the sample entropy, a measure of combinatorial coverage and chaos, decreases in further washes, clearly delineating that sequences that are increasingly strong binders to the substrate are indeed more related to each other. The trend should also imply that sequences in the eluate must then be even closely related and that eluate entropy and combinatorial coverage must be less than wash-3. However, surprisingly, the entropy, and therefore combinatorial coverage of the eluate is much higher than the weakest wash (More details in supplementary S3). The supposed strongest binders specific to the same substrate are more diverse and unrelated to each other than the weakest binders to the same substrate. Such an observation seemingly turns our previous assumption that strong binders to a specific substrate must be more alike than weak binders on its head. A word of caution is that the previous assumption does indeed hold, but only if the binding mechanism is the same. Following the previously established phage-display procedure, the commonly used generic tween 20/80 detergent didn’t categorically attack specific binding modes and mechanisms. Moreover, many eluted peptides had smaller but significant populations represented in the previous washes due to experimental error. The effect of this ‘information leakage’ may be mitigated by transforming the survival affinity into the ‘center-of-abundance-mass’ metric, as shown previously. Models that use the latter metric to regress or classify upon may be more successful at reconciling the trends observed in the training, test, and benchmark datasets.

## Conclusions and Future steps

We have developed a universal first step in high-throughput peptide selection approach to obtain solid-binding peptides in high-throughput and identified more than 2 Million different MoS_2_ binding peptides. Through this paper, we disseminate the first and largest ever solid binding peptide dataset labeled with survival probability scores for binding activity on Molybdenum Disulfide substrate, selected via a high throughput protocol that we adapt into GEPI selection. The dataset is cleaned, conditioned, and analyzed to guide the next stage in high throughput deep directed evolution experiments with functional scoring. Moreover, we also disseminate a carefully selected benchmark GEPI dataset for affinity to MoS_2_, characterized and labeled in terms of relative surface coverage under Fluorescent microscopy as well as phage retention from spectrophotometer absorbance. In addition to this, having a large number of experimentally selected peptides enables in-depth exploration of the sequence space. It allows Machine-Learning assisted designing of superior 2nd generation peptides with desirable properties.

The efficacy of previously and newly developed machine learning models to characterize the dataset was suboptimal due to the underlying assumption adopted from biology that denaturing the peptides is enough to disrupt the binding events. Just because a sequence is present in greater likelihood in the eluate and less in the washes does not make the sequence a strong binder to the substrate per se, but rather that the generic tween detergents are not disrupting the binding mechanism of that particular sequence. Denaturation is not enough. We conclude that the eluate peptides bind with various mechanisms not targeted by the tween detergent, necessitating a binding-mechanism targeted approach. Explicitly using different washing solutions that target separate substrate-phage interaction mechanisms such as hydrophobicity, aromatic-interaction, amino-acid competition, etc., will be crucial in further resolving the factors that govern binding interactions at the bio/nano interface. Moreover, because we see a significant number of peptides that appeared in washes also turn up in the eluate, refinement and optimization of the eluate via amplification steps coupled with next-generation sequencing protocol are warranted.

The conclusion necessitates a multiplexed approach with a greater number of intermediate panning and amplification steps to fully characterize the washes in terms of functional scores that indicate binding affinity. Information entropy, which measures diversity or the combinatorial coverage, is a good measure of the sample’s reliability in representing the population’s distribution. The entropy analysis outlined above is a simple yet successful method that can guide successive iterations (Fig 1 shows such an iterative process) of the deep directed evolution experiment that we have introduced.

Additionally, amplifying the eluted peptides multiple times and repeating the experiment seems like the natural next step to experimentally segregate the sequence space along differences in binding mechanisms. Such a new and improved experimental procedure, combined with the data exploration and quality control methods described in the current work, promises to generate high-quality massive datasets for a thorough exploration of evolutionary dynamics. It will enable highly efficient and rich directed evolution experiments to select novel proteins and peptides for practical implementations in technology and medicine and enrich our understanding of nature’s design process.

## Acknowledgments

The research is financially supported (DTY, SSR, JLR, JFL, ONB, MS) by National Science Foundation (NSF) through the DMREF program (via Materials Genome Initiative) under grant numbers DMREF DMR# 1629071, 1848911, and 1922020. We thank by Jason J. Stephany and Douglas Fowler for technical help in NextGen selection processes and the use of their facilities (UW Genome Sciences), and discussions in machine intelligence and guidance by Kevin Jamieson (Allen School of Computer Science and Engineering, University of Washington).

## Supplementary Information

### S1: Conventional Directed Evolution for Benchmark data Generation

The 12-mer Phage display (PhD) library is used to select peptide sequences fused to the minor coat protein (pIII) against MoS_2_ flakes as described above. The obtain enough number of clones for subsequent rounds, eluted phages are transferred to an early log phase E. coli ER2738 culture (∼OD: 0.4) and amplified for 4 hours. The cell pellet is obtained by centrifugation and purified by polyethylene glycol (PEG) precipitation according to the manufacturer’s instructions. Purified phages are obtained in PC buffer volume pH 7.4 with a final volume of 200 μl. For the individual phage isolation, eluate pools are streaked on agar plates and individual colonies were picked. Amplification is performed in the early log phase E. coli ER2738 culture for 4 hours followed by polyethylene glycol (PEG) precipitation.

The single-stranded DNA of selected phage plaques are isolated by a QIAprep Spin M13 Kit (Qiagen, Valencia, CA) and amplified via PCR in the presence of dye-labeled terminators (Big dye terminator v3.1, Applied Biosystems, Carlsbad, CA). PCR products are purified by Sephadex G-50 column precipitation. A 96 gIII primer (5′-OH CCC TC TAGTTA GCG TAA CG-3′) is used for the amplification of ssDNA. The selected sequences of DNA from clones are analyzed by an Applied Biosystems 310 Avant DNA analyzer. After sequencing, the isolated clones are then tested individually for their relative binding affinities to MoS_2_. To measure the relative binding affinity, two different assays are used.

Firstly, the individual colonies are mixed with 5 mg of MoS_2_ flakes dispersed in 1 mL of PC Buffer (pH 7.4), containing 0.02% Tween 20 detergent and incubated for 3 hours on a rotator at room temperature. After washing off loosely bound phages from the MoS_2_ surface using 1 mL of PC Buffer (pH 7.4), containing 0.1% Tween 20 detergent, the bound clones are fluorescently labeled using anti-M13 antibodies. The samples are characterized by quantitative fluorescent microscopy, employing a Nikon Eclipse TE-2000U fluorescent microscope (Nikon, Melville, NY, USA) equipped with a Hamamatsu ORCA-ER cooled CCD camera (Hamamatsu, Bridgewater, NJ, USA), imaged using a FITC filter (exciter 460–500 nm, dichroic 505 nm, emitter 510–560 nm) and MetaMorph imaging system (Universal Imaging, West Chester, PA, USA). Finally, the binding affinity for each peptide is determined by calculating the ratio of total fluorescent intensity over total surface area of the MoS_2_ flakes (n=10). Secondly, for spectrophotometry based analysis, number of bound phages are determined by analyzing the broad optical absorption peak located from 260 nm to 280 nm, with a slight maximum at 269 nm which reflects the nucleotide content of the particular phage, a molar extinction coefficient (9.006×103 M_-1_cm_-1_) where the genome size of M13KE is taken as 7222 base-pairs. Prior to MoS_2_ incubation, the initial concentration of each clone is calculated. Next, the relative binding affinities are calculated from the measured absorption intensities of depleted (unbound) phage solutions as a percentage of the original absorption intensity from starting solutions. Each experimental set is performed in triplicate.

A total of 144 colonies are isolated and sequenced from each eluate (36 colonies after each biopanning) which yielded a total of 96 unique sequences. As shown in Fig 2, the binding coverage of selected peptides ranges from 20% to 90% demonstrating the presence of strong, moderate, and weak binders. The fluorescent signal is obtained through immunolabeling by fluorescently labeled anti-M13 antibodies and therefore the total intensity is a function of the number of antibodies on the surface. It is important to note that depending on the average number of antibodies on the phage surface, the total intensity can vary significantly. On the other hand, in spectrophotometric method, absorption at maximum at 269 nm is a direct measurement for the total genomic DNA content in a phage solution which provides a direct quantification for phage concentration present on the surface. As shown in Fig 2, the results of each assay have about 70 percent overlap, with 4 exceptions (clone 1, 21, 34, 54). One possible explanation for the discrepancy between the two assays, in particular, clone# 1, 21, 34, 54, could be the repeated centrifugation steps where the phages can be trapped between the pelleted flakes and causes false negatives in fluorescent imaging. Moreover, in spectrophotometric characterization, the incomplete sedimentation of tiny MoS_2_ particles could cause false positives with increased UV absorption.

**S1 Fig.**
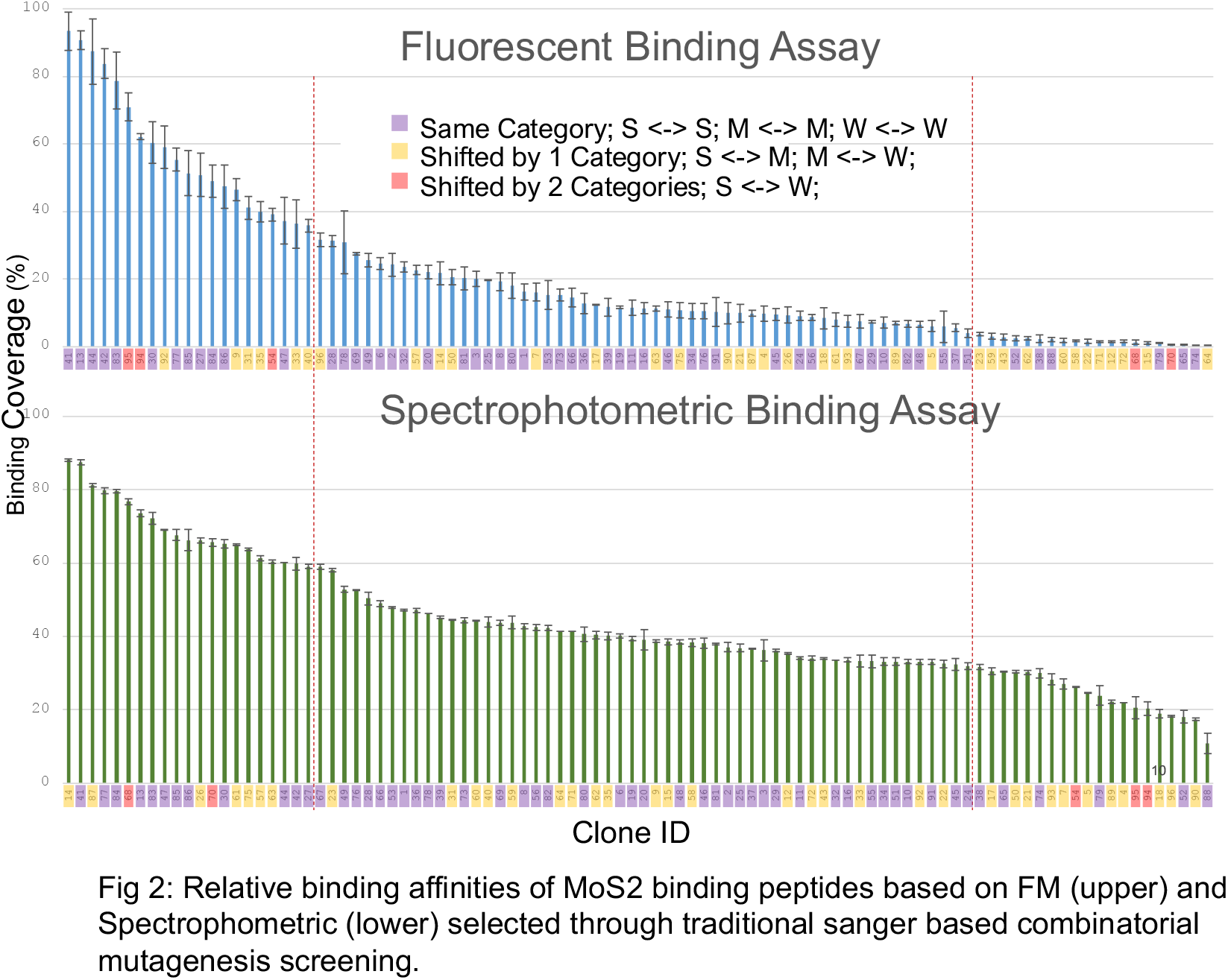
**Fluorescent Microscopy and Spectrophotometry characterization of phage retention for sequences obtained through conventional directed evolution and sanger sequencing**.

### S2: Predictive Modelling

In previous work, it was shown that the binding affinity of a peptide of interest is proportional to the sum of the Global Alignment scores to a group of highly functional peptides. Like the novel approach described here, this functional scoring, referred to as the Total Similarity Score (TSS) trend was compared with the experimental binding trend generated by the fluorescence microscopy procedure described above. Traditional models previously generated in GEMSEC were applied in addition to expanded applications of the TSS concept to diversify our analysis.

#### Traditional Similarity Analysis/Iterative Alignment

Similarity Analysis, or Iterative Alignment, is the first computational scoring method that was developed by GEMSEC, being benchmarked on quartz binding peptides in 2007. Essentially, Similarity Analysis, or Iterative Alignment, optimizes a similarity matrix (distance matrix between amino acids) that ensures the most active (binding, mineralization etc.) peptides are more similar to each other than they are to the least active peptides. The Total Similarity can be explicitly stated as the averaged Needleman-Wunsch alignment score of a group of sequences A to another group B, where the Total Similarity Score of A to B is written TSS_A-B_. The internal similarity of the highly active peptides is captured by the Total Similarity Score Strong-Strong (TSS_S-S_). External similarity of the highly active to the least active is referred to as the Total Similarity Score Strong-Weak (TSS_S-W_). Random changes to the similarity matrix that increase the TSS_S-S_ and decrease TSS_S-W_ are considered ‘beneficial changes’ and are done until specified by the experimenter. In general, the sequences should become more and more related in sequence as the peptides become more similar in function, leading to a characteristic trend demonstrated by the quartz application (S3 Fig). The seed matrix most commonly used, PAM250 was derived in the seventies by leveraging how mutations affected the function of proteins between closely related species and served as a successful starting point for trained matrices in the original work. In order to generate the characteristic trend of internal similarity to confirm the directed functionality of the dataset, PAM250 matrices were trained using 1000 highly active peptides (highest survival affinity) and 1000 less active peptides (lowest survival affinity) from each biological replicate, including a combined set ensuring consistency of survival affinity across at least 2-3 sets. These matrices were then used to score 5 groups of sequences taken from the three 2.5 million data-points (Strong, Less Strong, Medium, Less Weak and Weak). The 96 Sanger MoS_2_ binding peptides were used to benchmark the matrices trained on the larger dataset with affinity for the same material by classifying each sequence by their affinity (Strong, Medium, Weak). Although the trend of internal similarity tended to decrease among all the bar charts generated (Supplementary Section, S3 Fig), the matrices from all datasets except the combined set struggled to place the benchmark sequences in their correct affinity class (Supplementary Section, S3 Fig).

#### Machine-learning on Total Similarity Scores

The method referred to as ML on TSS (Machine Learning on Total Similarity Scores) is best characterized as a multiple regression on eight independent Total Similarity Scores featurized by 550+ physiochemical properties of a peptide set towards prediction of their survival affinity downloaded from a database called the Amino Acid Index, which contains hundreds of curated properties that have been experimentally measured. A schematic is shown in S4 Fig. The peptides used as the highly functional group were those sequences with the highest survival affinity values. The distances between AAs were derived by performing Principal Component Analysis (PCA) on each group of properties to generate ∼20 orthogonal variables by which each amino acid could be described by its Cartesian distance to the others. In the next step, the similarity matrices were populated by the appropriate distances from one AA to another, creating a symmetric matrix. The sequences were scored by calculating their TSS with respect to the most active peptides (the strongest binders with the highest survival affinity) using each similarity matrix, generating 8 scores per peptide. The experiment performed herein was done to explore the effect of the copy number (total number of copies of a particular peptide present during the wash process) on the prediction accuracy. 9 datasets were created that were distinguished by the minimum copy number of peptides that were identified in each biological replicate. The regression was overall able to predict the survival affinities of other sets and themselves with high accuracy (as low of a mean square error as ∼7 for the most stringent count discrimination; S4 Fig). As the count discrimination increases, the mean square error decreases, indicating the data containing the most information about the overall trend is present in the sequences having more copies in the selection process.

#### Total Similarity Score – Convolutional Neural Network

To deepen the learning space of the methods while keeping to the cost function known to describe peptide activity, a constrained convolutional neural network was generated that ensures the parameters necessitated by similarity matrices (symmetric matrix) and position-specific frequency matrices (columns per position must sum to 1). Neural networks have been used to classify and quantify the affinity of smaller datasets of thousands of peptides with reasonable success, indicating a large amount of learning space is required to capture the informatic trend. The TSS-CNN method can be described as a simple constrained neural-network trained to generate a trend of affinity scores that correspond to the -x^3^ curve, the trend always generated by traditional similarity analysis. A schematic of the process is given by S5 Fig. The strength of this method is in its ability to generate the full representative space of how peptides bind to MoS_2_ (or the directed function of the dataset) by optimizing the locations and is to identify two matrices of size (12×20) and one of (20×20) are randomly initialized and their constraints applied. Next, the TSS to the frequency matrix representing the strongest binders is calculated per peptide by multiplying the frequency of occurrence of all AAs in the column corresponding to the position of the peptide of interest by the mutability of the AA in that position. The process is repeated across all positions until the entire peptide has been effectively matched with the entire group of high survival affinity peptides (represented by the frequency matrix). The values calculated per position and across the peptide are summed to create a single TSS for the peptide. When applied to a set with directed functionality (i.e. binding to MoS_2_), a distribution is generated. Changes to the similarity matrix and frequency matrices were kept dependent on whether they increased the nonlinear correlation of these TSS values with an evenly spaced distribution of values on the -x^3^ trend. These matrices were expected to be representative of the peptide binding trend defined by the surface. The first iteration of the CNN failed to predict a trend in affinity for the benchmark peptides. Compared to the fluorescence (green line, S5 Fig) and spectrophotometric (orange line, S5 Fig) trends, the TSS trend (blue lines, S5 Fig) is unable to group binders into semi-quantitative classes of strong, medium, and weak. Although the sum of all the frequencies per position in the Position Specific Frequency Matrix summed to 1, the fifth position blew its values out of proportion with the rest, likely attributable to a wide diversity of amino acids giving a wide range of functions in this position. In the updated method new constraints will be applied such that the values must remain within a certain range.

#### Genetic Algorithm

The purpose of this method is to diversify the approach beyond machine learning and leverage the size of the dataset to identify changes in sequence that would change the function of the benchmark peptides in a controllable manner. This method focused on identifying mutational changes in peptides similar in sequence to the original 96 MoS_2_binding peptides with the goal of predictively designing the affinity of new MoS_2_ binders. A schematic of the full process is provided by S6 Fig. Once the full list of similar peptides had been identified, they were filtered by the consistency of their survival affinity across the 3 biological replicates, to ensure the data was the most accurate. Further, only sequences with at least 8 total copies during the selection process were included in the final lists. After the cleaning process was completed, the differences in binding affinity between sequences only 1 AA apart was calculated and tracked along with the mutation and the position where it occurred. These changes in amino acid sequence were identified by first finding peptides within each of the 3 biological replicates that contained at least 7 amino acid (AA) matches (having the same amino acid in the same position) with the 96 Sanger peptides that were selected for affinity to MoS_2_. This process was repeated 5 times to ensure similarity to the original sequences was maintained and to capture a sizeable portion of the dataset. This information was graphed in a 3D matrix (x = Initial AA, y = Final AA, z = Position) where the color indicates the change in affinity associated with the mutation (S6 Fig). The standard deviation of each affinity change was also graphed to ensure mutations tested had a consistent effect across peptides. Overall, the mutations affecting the survival affinity of the peptides were the cysteine to alanine mutations in position 4 (fourth amino acid from the N terminus) which was associated with a dramatic loss of function, while the largest gain-of-function mutations were centered around the mutation of cysteine to leucine and tyrosine to asparagine. It is reasonable to attribute the gain-of-function results to the affinity of asparagine (positively charged) for the negatively charged MoS_2_ surface. Further, the loss of a highly negatively charged group (cysteine) to a very hydrophobic chain (leucine) may be indicative of a larger trend of hydrophobicity lending weight to the strength of the interaction.

**S2 Fig.**
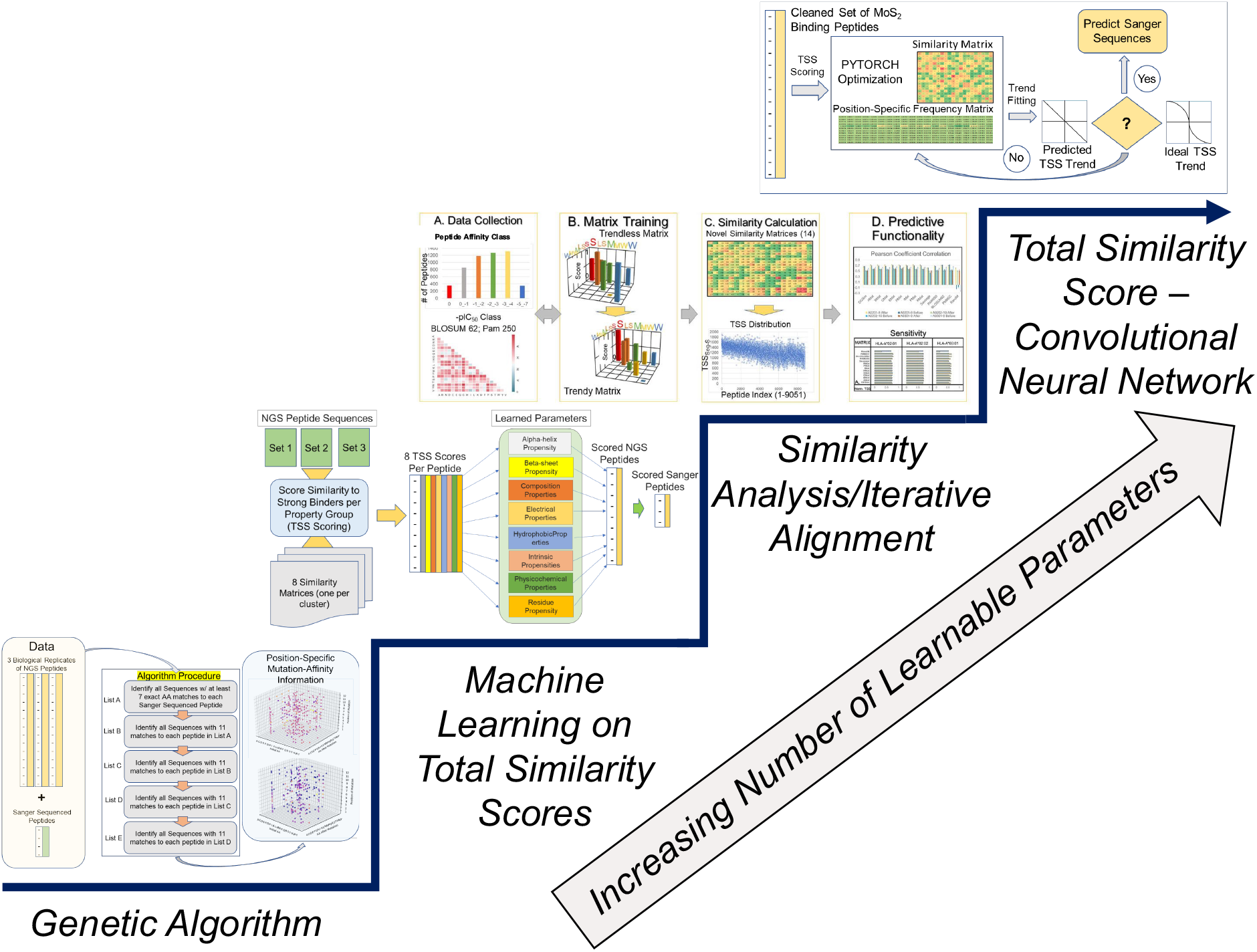
**Increasingly complex mathematical models for preliminary predictive analysis on the dataset**.

**S3 Fig.**
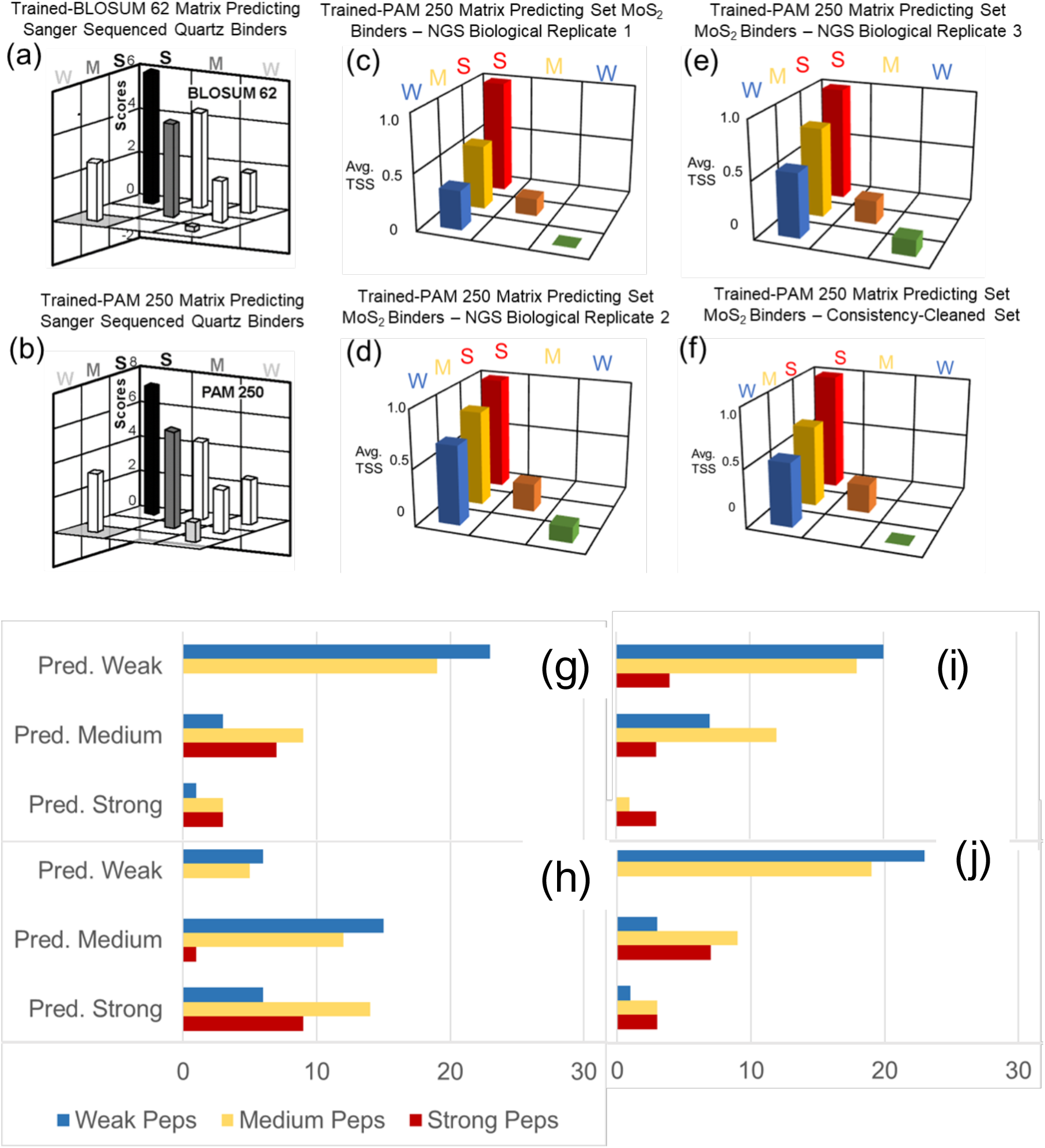
Traditional Similarity Analysis and Iterative Alignment process.

**S4 Fig.**
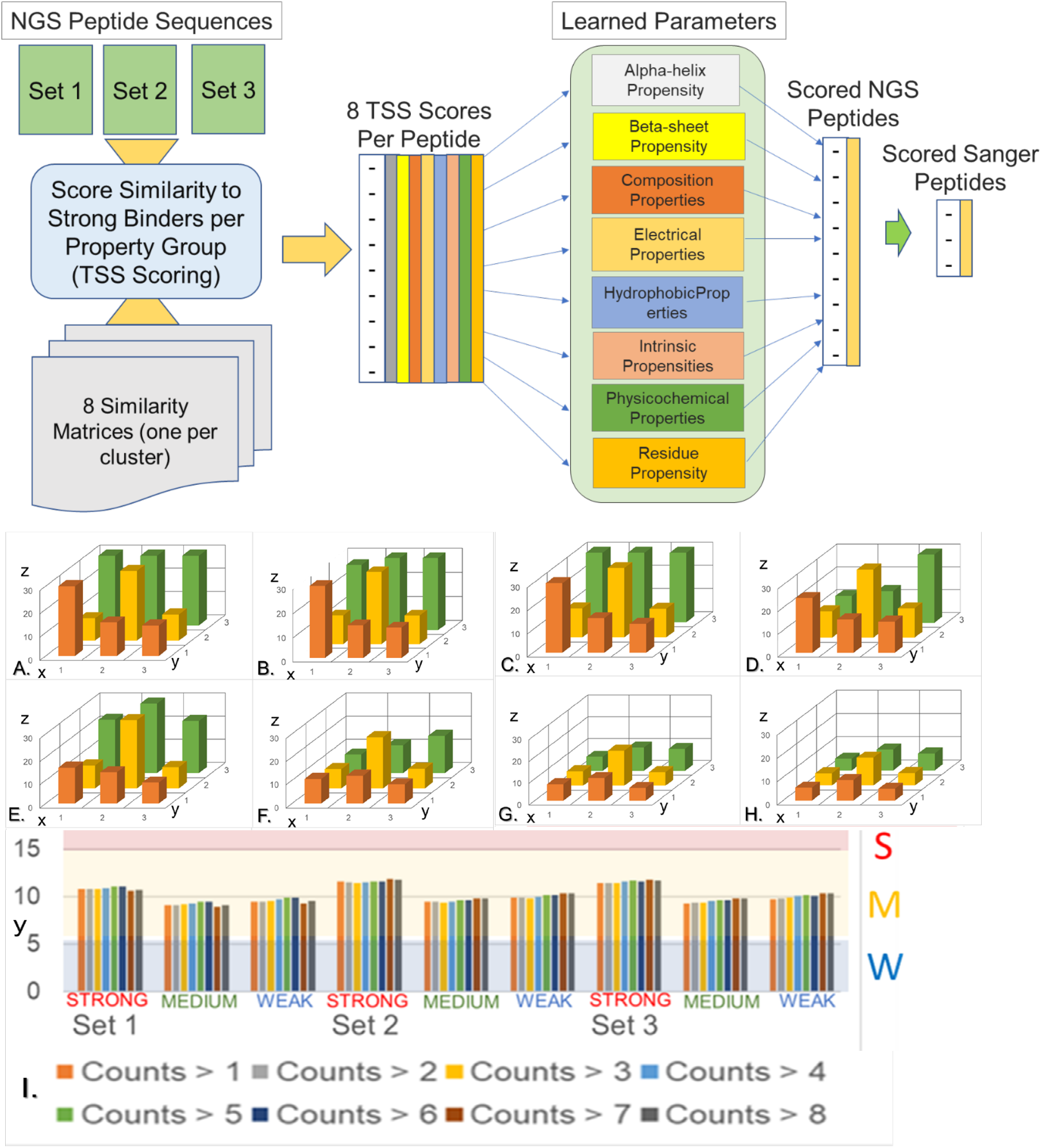
Machine Learning on TSS Scores.

**S5 Fig.**
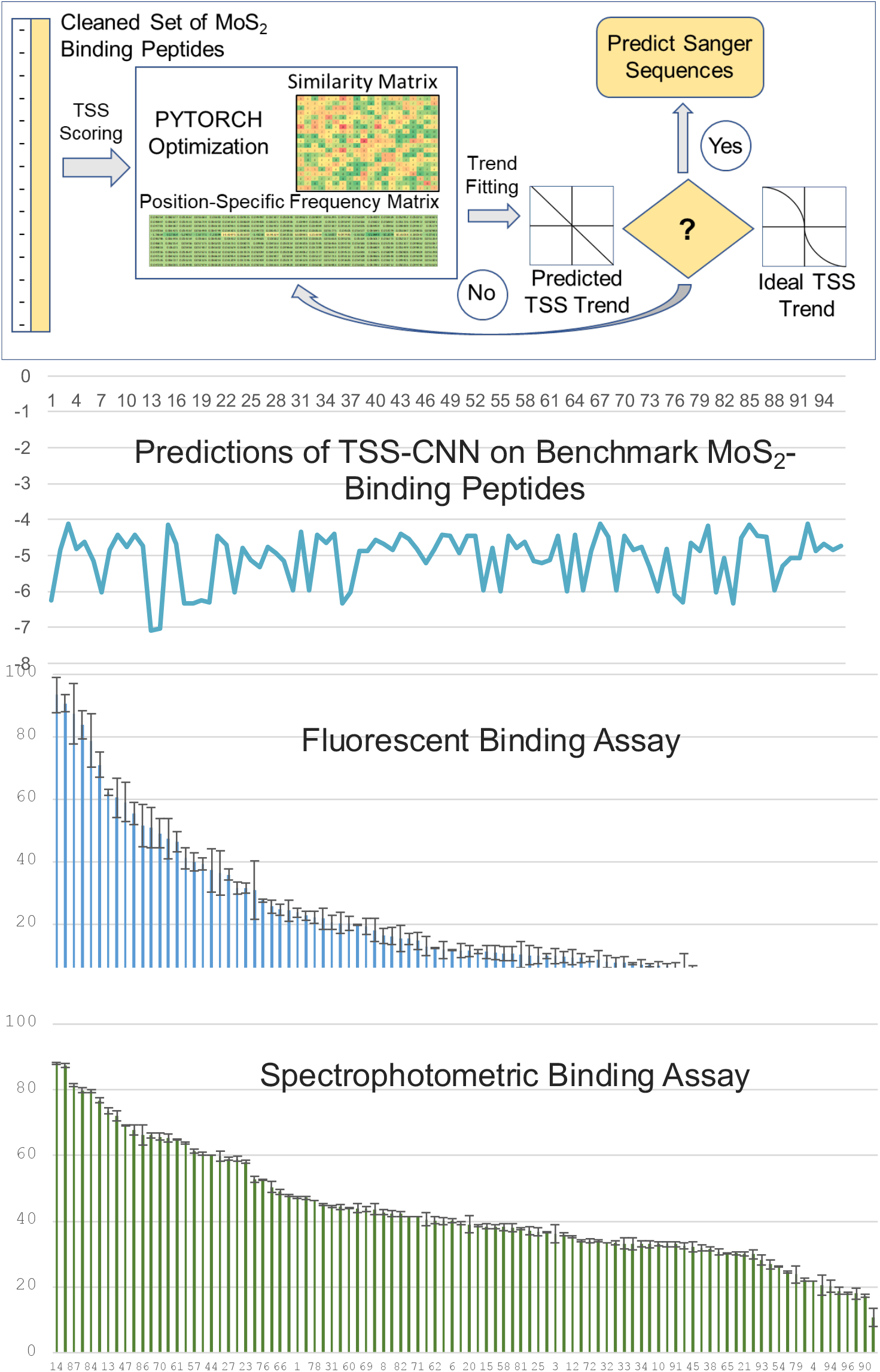
TSS-CNN: Applying a simple feed forward neural network on the TSS scores.

**S6 Fig.**
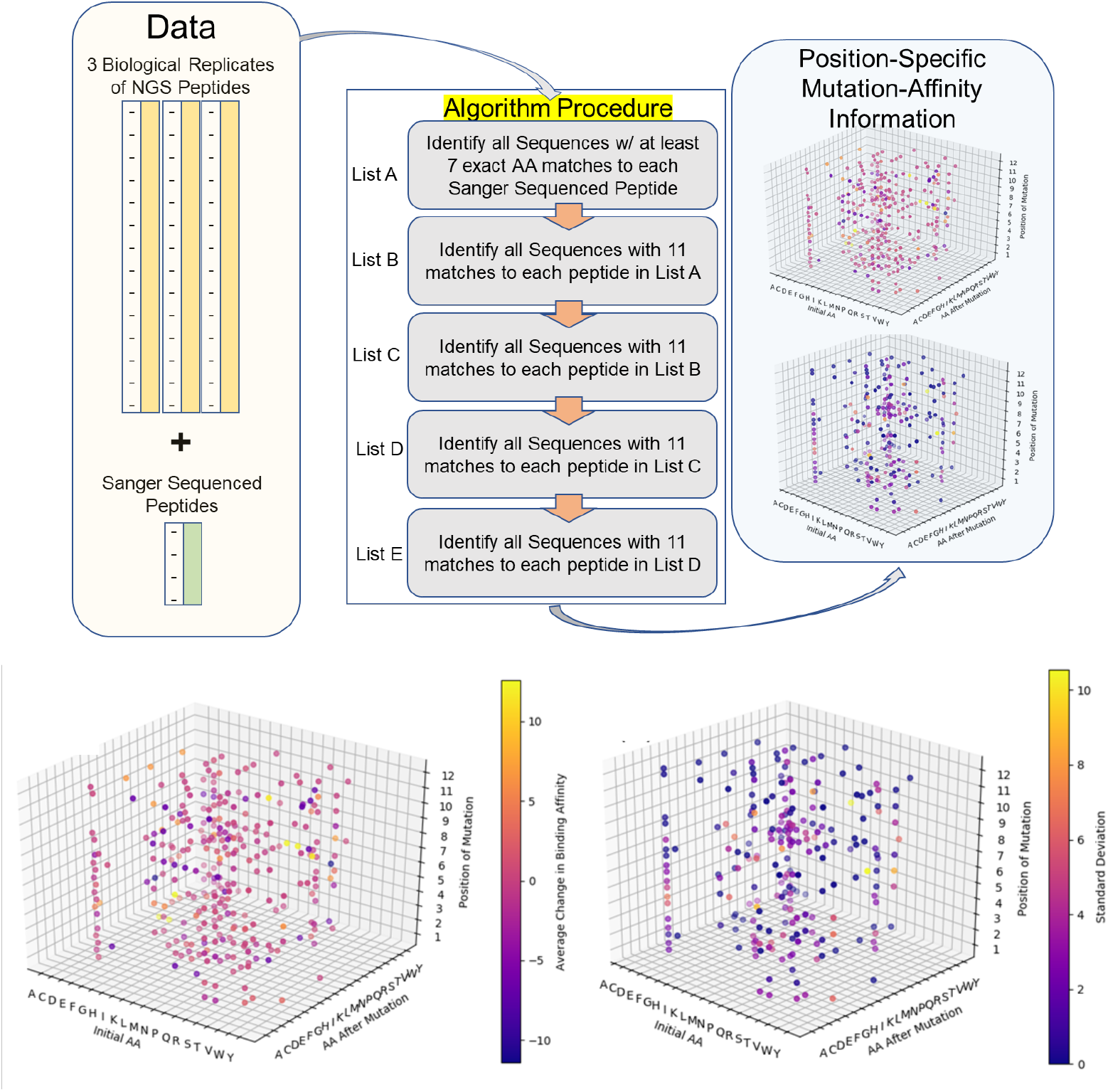
Genetic Algorithm Schematic and results of evolutionarily conserved mutations as described in S2.

### S3: Assessment of Diversity by Random Sampling of Phage Pools

To further analyze the diversity of the sequences, we sample 2, then 20, then 200 (with replacement) and continue sampling until we reach 2 million DNA sequences from each wash and eluate phage pool and calculate the number of unique sequences per draw. If the number of unique sequences (by extension, diversity) increases upon continuously sampling more sequences from the same wash/eluate, then the minimum sample size has not been reached that would satisfactorily describe the behavior of the theoretical population in that phage pool. Increasing number of unique sequences upon larger samplings from the same wash/eluate would mean that there are still more combinations of nucleotides available that would display the same behaviors. While a sigmoidal nature of the trend, upon continuously increasing sample size from the same wash/eluate would indicate that we are approaching the sample size that satisfactorily represents all/most possible combinations of nucleotides. In short, increasing trend of entropy indicates that sample size is not large enough to fully describe the combinatorial space of the wash/eluate while a sigmoidal entropy trend indicates the opposite. As seen in the figure below, the uniqueness trends are exponentially increasing for Wash 1 and Wash 2 in all biological replicates of the experiment, while it is stabilizing for Wash 3. The results prove that indeed Washes 1 and 2 do not yet have a representative combinatorial coverage of sequence space to describe weak binders, while Wash 3 is approaching a sample size which would possess a representative combinatorial coverage of sequences to describe the strong binders that bind with a mechanism that is disrupted by the tween detergents. The eluate deviates from this trend and entropy increases as we randomly sampled more and more sequences from the eluate, indicating that the supposed strong binders are indeed binding with multiple mechanisms that necessitate a more targeted approach to directed evolution and NGS protocol than that used in this study. Moreover, because we see a significant representation of peptides that appeared in washes also turn up in the eluate, and since the input library was not explicitly sequenced, a refinement and optimization of the current directed evolution and next generation sequencing protocol is warranted.

**S7 Fig.**
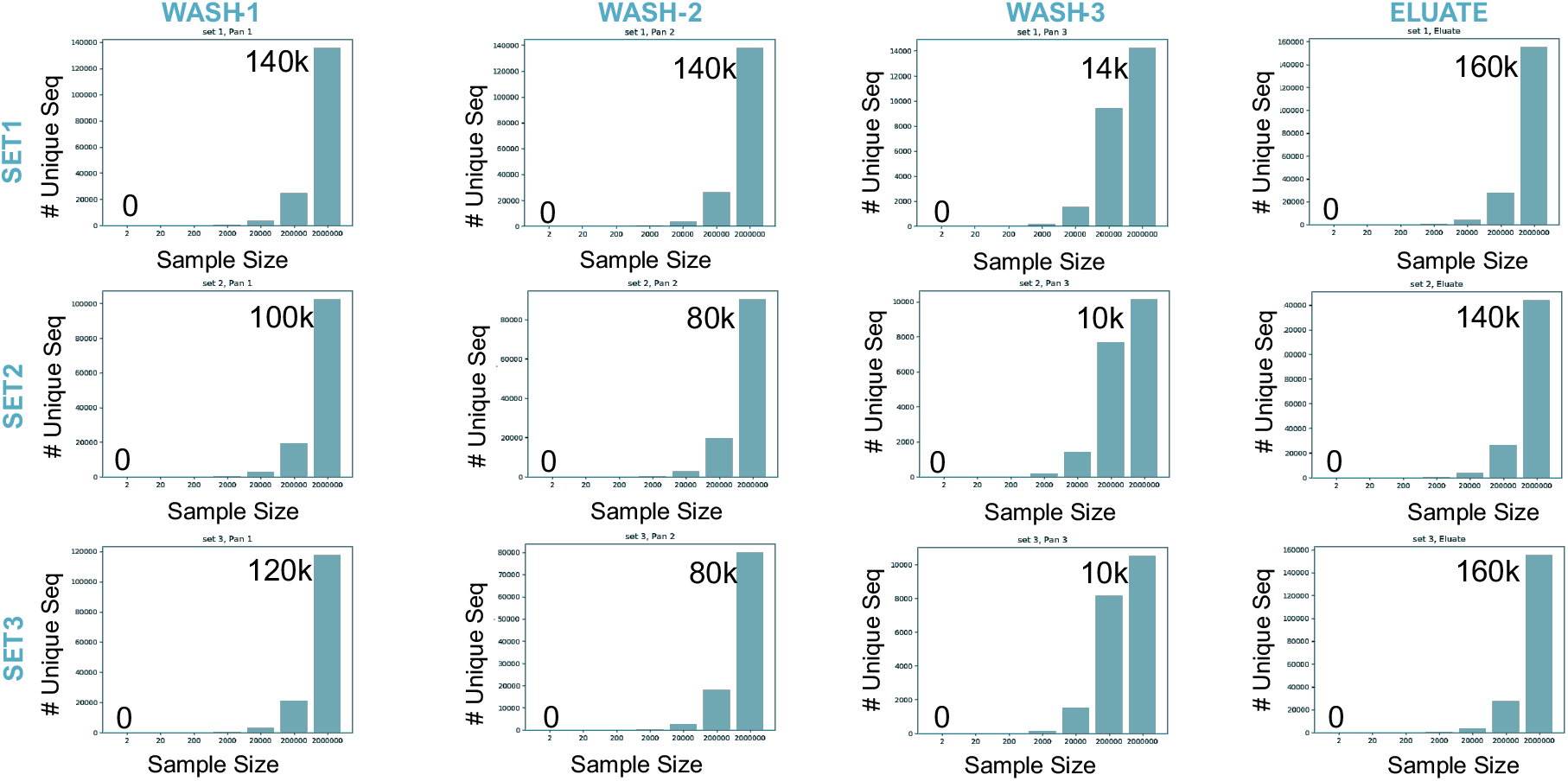
assessment of diversity and sample size requirements in phage pools.

